# Hinge Influences in Murine IgG Binding to *Cryptococcus neoformans* Capsule

**DOI:** 10.1101/2021.01.18.427174

**Authors:** Diane Sthefany Lima de Oliveira, Verenice Paredes, Adrielle Veloso Caixeta, Nicole Moreira Henriques, Maggie P. Wear, Patrícia Albuquerque, Maria Sueli Soares Felipe, Arturo Casadevall, André Moraes Nicola

## Abstract

Decades of studies on antibody structure led to the tenet that the V region binds antigens while the C region interacts with immune effectors. In some antibodies, however, the C region affects affinity and/or specificity for the antigen. One such case is that of the 3E5 antibodies, a family of monoclonal murine IgGs in which the mIgG3 isotype has different fine specificity to the *Cryptococcus neoformans* capsule polysaccharide than the other mIgG isotypes. Our group serendipitously found another pair of mIgG1/mIgG3 antibodies based on the 2H1 hybridoma to the *C. neoformans* capsule that recapitulated the differences observed with 3E5. In this work, we report the molecular basis of the constant domain effects on antigen binding using recombinant antibodies. As with 3E5, immunofluorescence experiments show a punctate pattern for 2H1-mIgG3 and an annular pattern for 2H1-mIgG1. Also as observed with 3E5, 2H1-mIgG3 bound on ELISA to both acetylated and non-acetylated capsular polysaccharide, whereas 2H1-mIgG1 only bound well to the acetylated form, consistent with differences in fine specificity. In engineering hybrid mIgG1/mIgG3 antibodies, we found that switching the 2H1-mIgG3 hinge for its mIgG1 counterpart changed the immunofluorescence pattern to annular, but a 2H1-mIgG1 antibody with a mIgG3 hinge still had an annular pattern. The hinge is thus necessary but not sufficient for these changes in binding to the antigen. This important role for the constant region in binding of antibodies to the antigen could affect the design of therapeutic antibodies and our understanding of their function in immunity.

**Key points:**

- Key point 1- 2H1 antibodies recapitulate differences between mIgG isotypes observed with 3E5.
- Key point 2 – The hinge region is necessary but not sufficient for these differences.
- Key point 3 - The antibody constant region can also play a role in mIgG binding to antigen.

## Introduction

Since their discovery in 1890 (1), antibodies have been one of the most studied and most useful biomolecules. Their enormous versatility for both the immune response in animals and for biotechnology is in great part due to the fact that they are bifunctional, with a highly variable end that binds to antigens with exquisite specificity and a constant end that is bound by receptors in immune effector cells. More than half a century of studies on the structure and function of antibodies, starting with seminal work by Porter and Edelman (2–4), have shown that the prototypical IgG molecule is a glycosylated heterotetramer composed of two Fab domains connected to an Fc domain by a flexible hinge (5). The heavy and light chain monomers form complete immunoglobulin (Ig) molecules that can be classified into five types: IgG, IgM, IgA, IgE, and IgD (6–8). Among the classes, the IgG isotypes are the most studied because they engage specific Fc receptors and their effectiveness in neutralizing pathogens and toxins, opsonizing targets for phagocytosis and antibody-dependent cellular cytotoxicity, activating complement and modulating the inflammatory response. Due to their numerous functions, IgG antibodies naturally play crucial roles both in health and in disease (9), are the basis for dozens of drugs used to treat illnesses (10, 11) and are widely used as diagnostic and biotechnological tools.

Until recently, the most widely accepted model for antibody structure is one in which specificity and affinity for the antigen is determined solely by the variable (V) region, whereas the constant (C) region determines antibody effector functions and class/isotype. This concept emerged from the understanding that V and C regions functioned independently without affecting each other’s activity. Since the 1980’s, a series of antibody studies have questioned this classical definition, instead suggesting intramolecular or allosteric cooperativity between antibody C and V regions (12–20). Evidence for this comes mostly from experiments in which different IgG isotypes with identical variable regions had differences in antigen binding. Such evidence suggests that the structure-function model of the antibody molecule needs revision.

Studies from our group have demonstrated the impact of the C region on murine IgG (mIgG) antigen affinity and specificity using *Cryptococcus neoformans* as a model with different antibody sets (14–17, 21). The most studied set of identical V region antibodies is the 3E5 mIgG1, mIgG2a, mIgG2b and mIgG3 hybridoma mAbs that were generated by isolating spontaneous isotype-switched hybridoma variants by cell culture techniques (22). The specificity, affinity and structure of these and other antibodies has been extensively studied by immunofluorescence (16), ELISA (22), phagocytosis assay (23), isothermal titration calorimetry (15), surface plasmon resonance (24), molecular modeling (25), circular dichroism (26), nuclear magnetic resonance, fluorescence emission spectroscopy, molecular dynamics simulations (27), small-angle X-ray scattering and X-ray crystallography (25). Immunofluorescence experiments using 3E5 have demonstrated that 3E5-mIgG3 binds to the *C. neoformans* capsule in a punctate pattern, which differs from the annular pattern observed with other mIgG isotypes. This difference in binding pattern correlates with antibody protective efficacy (28, 29). ELISA with regular (acetylated) and de-O-acetylated capsular polysaccharide demonstrated that 3E5-mIgG3 had a much higher affinity for the de-O-acetylated polysaccharide when compared with the other mIgG isotypes, whereas all bound acetylated GXM with similar affinities. Both observations indicate an influence of the C region in antigen binding in 3E5 antibodies. Strikingly, this paralleled a difference in passive immunization experiments with mice: mIgG3 is non-protective or disease enhancing, whereas the other mIgG isotypes protected mice infected with *C. neoformans* (17, 22). Circular dichroism further demonstrated that mIgG3 antibodies suffer allosteric changes to their structure upon exposure to antigen (13, 26). These and other results suggest that there is a strong correlation between the protective mechanism of antibodies and antigen binding. However, the strength of coupling between V and C regions appears to vary for different combinations such that some are more conducive to structural communication than others (12).

Changing the C region while maintaining the same V region is a crucial step in the biotechnological development of therapeutic and diagnostic antibodies. It is also very important for an effective immune response in animals, which switch antibody isotypes depending on the type of antigen present. Given that C region changes might not only affect the paratope but can transform a protective antibody into a disease-enhancing one, clarifying these effects is important both for our understanding of the immune system and also for the safe and effective development of therapeutic antibodies. While carrying out studies on the mIgG3 receptor (23) we generated recombinant mIgG1/mIgG3 antibodies to the *C. neoformans* capsule that serendipitously showed the same difference in immunofluorescence pattern as the 3E5 antibodies. These recombinant antibodies are based on the V region sequences of the 2H1 hybridoma, originally an mIgG1 mAb against the *C. neoformans* capsule that differs from 3E5 in four positions on the variable kappa (VK) sequence and eight residues on the VH domain. In this work, we followed up on that observation by engineering these mIgG1/mIgG3 antibodies to switch CH1 and hinge regions between the mIgG1 and mIgG3 isotypes to understand the structural mechanism that leads to their differences in specificity and affinity to their antigen.

## Materials and Methods

### Fungal strains and culture

We used the wild type H99 (serotype A, ATCC^®^ 208821™) and the Cas1 knockout mutant (*cas1Δ*) strains of *C. neoformans*. The Cas1 gene encodes an O-acetyltransferase that is necessary for acetylation of capsular polysaccharides (30). Yeast cells were maintained in Sabouraud agar plates and grown in Sabouraud dextrose broth (Difco) or minimal medium (0.3% glucose, 13 mM glycine, 29.4 mM KH_2_PO_4_, 10 mM MgSO_4_, 3 μM thiamine, pH 5.5) at 30 °C at 150 rpm for 2 days.

### Cell lines

The following cell lines were used: J774.16 – murine macrophage-like cells; CHO dhFr−/−(ATCC^®^ CRL-9076) – Chinese hamster ovary cells lacking dihydrofolate reductase; NS0 (ATCC^®^ PTA-3570) – murine myeloma. J774.16 and CHO dhFr−/−cells were grown at 37°C and 5% CO_2_ in Dulbecco’s Modified Eagle’s Media (DMEM) supplemented with heat-inactivated 10% fetal bovine serum (from South American origin), 10% NCTC-109 Media (Thermo Fisher) and 1% non-essential amino acids (Thermo Fisher). CHO dhFr−/−cells were also supplemented with 1x sodium hypoxanthine and thymidine (HT) solution (Gibco). NS0 cells were maintained in the same supplemented medium and were grown in CD Hybridoma chemically defined medium (Thermo Fisher) in a 37°C incubator with 125 rpm shaking and 5% CO_2_. Adherent cells were grown on tissue culture treated plates (BD Falcon) and removed by treatment with 3-5 mL trypsin (Gibco) then pelleted by centrifugation at 300 × g for 10 min, and finally re-suspended at the appropriate cell density in DMEM.

### Exopolysaccharide (EPS) purification

H99 and *cas1Δ* cells were grown in minimal media for five days. Cells were removed by centrifugation and filtration, and the supernatants concentrated by ultrafiltration with a 100 kDa membrane (Millipore) (31). The viscous layer containing EPS was harvested and dialyzed against distilled water. The EPS was then lyophilized, weighted and dissolved in ultrapure water. H99 EPS was de-O-acetylated by alkaline hydrolysis in ammonium hydroxide (pH 11.25–11.50) overnight (17, 32).

### Nuclear magnetic resonance (NMR)

*C. neoformans* EPS was prepared as indicated above. After lyophilization, the < 100 kDa fraction was solubilized in deuterated water (D_2_O). 1D [^1^H] NMR (600 MHz) spectra were recorded on Bruker spectrometers equipped with Avance II console and triple resonance, TCI cryogenic probe with z-axis pulsed-field gradients at 30°C in D_2_O. [^1^H] NMR spectra were standardized against the residual solvent peak (internal D_6_DSS, δ = 0.00 ppm). All experiments were conducted with 64, 128, or 256 scans and an FID size of 16384 points. Standard Bruker pulse sequences were used to collect the 1D data (p3919gp and zggpw5). Data were processed in Topspin (Bruker version 3.5) by truncating the FID to 8192 points, using a squared cosine bell window function, and zero filling to 65536 points.

### Hybridoma antibodies

The *C. neoformans*-specific monoclonal antibodies used in these studies were originally isolated following immunization of mice with GXM conjugated to tetanus toxoid (33). 3E5 mAbs are mIgG1, mIgG2a, mIgG2b or mIgG3 (34), whereas mAbs 12A1 and 13F1 are mIgM (33–36).

### Recombinant antibodies - synthetic genes and cloning

Given that mAbs 2H1 (mIgG1) and 3E5 are the only antibodies to the *C. neoformans* capsule to have their structure solved by crystallography (25),(25), we decided to create recombinant mIgG1 and mIgG3 mAbs with identical 2H1 V regions (Figures 1A-B). To produce 2H1 and 2H1-hybrid mIgG1/mIgG3 recombinant antibodies, the VH and VL regions of 2H1 and 2H1-hybrid antibodies were codon-optimized using a proprietary algorithm and synthetized by GenScript and then cloned into the commercially available pFUSE vectors (InvivoGen). These vectors contain the C region sequences for murine (strain 129S) γ1 and γ3 heavy chains and κ light chain. The VL sequence for 2H1 was inserted into the pFUSE2-CLIg-mK vector to generate the 2H1-VL_pFUSE2-CLIg-mKvector [MT019334]. The 2H1 VH sequence was inserted into the pFUSE-CHIg-mG1 and pFUSE-CHIg-mG3 vectors to generate the pFUSE-CHIg-2H1-mIgG1 [MT019335], and the pFUSE-CHIg-2H1-mIgG3 [MT019336] vectors. The 2H1-hybrid antibodies were created by changing the CH1 or hinge sequences between 2H1-VH_pFUSE-CHIg-mG1 and pFUSE-CHIg-2H1-mIgG3, resulting in the following five hybrid vectors: 2H1-VH_pFUSE-CHIg-mG3-CH1a-1 [MT019339], 2H1-VH_pFUSE-CHIg-mG3-CH1b-1 [MT019340], 2H1-VH_pFUSE-CHIg-mG3-CH1c-1 [MT019341], 2H1-VH_pFUSE-CHIg-mG1-h-3 [MT019337] and 2H1-VH_pFUSE-CHIg-mG3-h-1 [MT019338]. All constructs were confirmed by digestion, PCR and Sanger sequencing before transfection.

**Figure 1.**
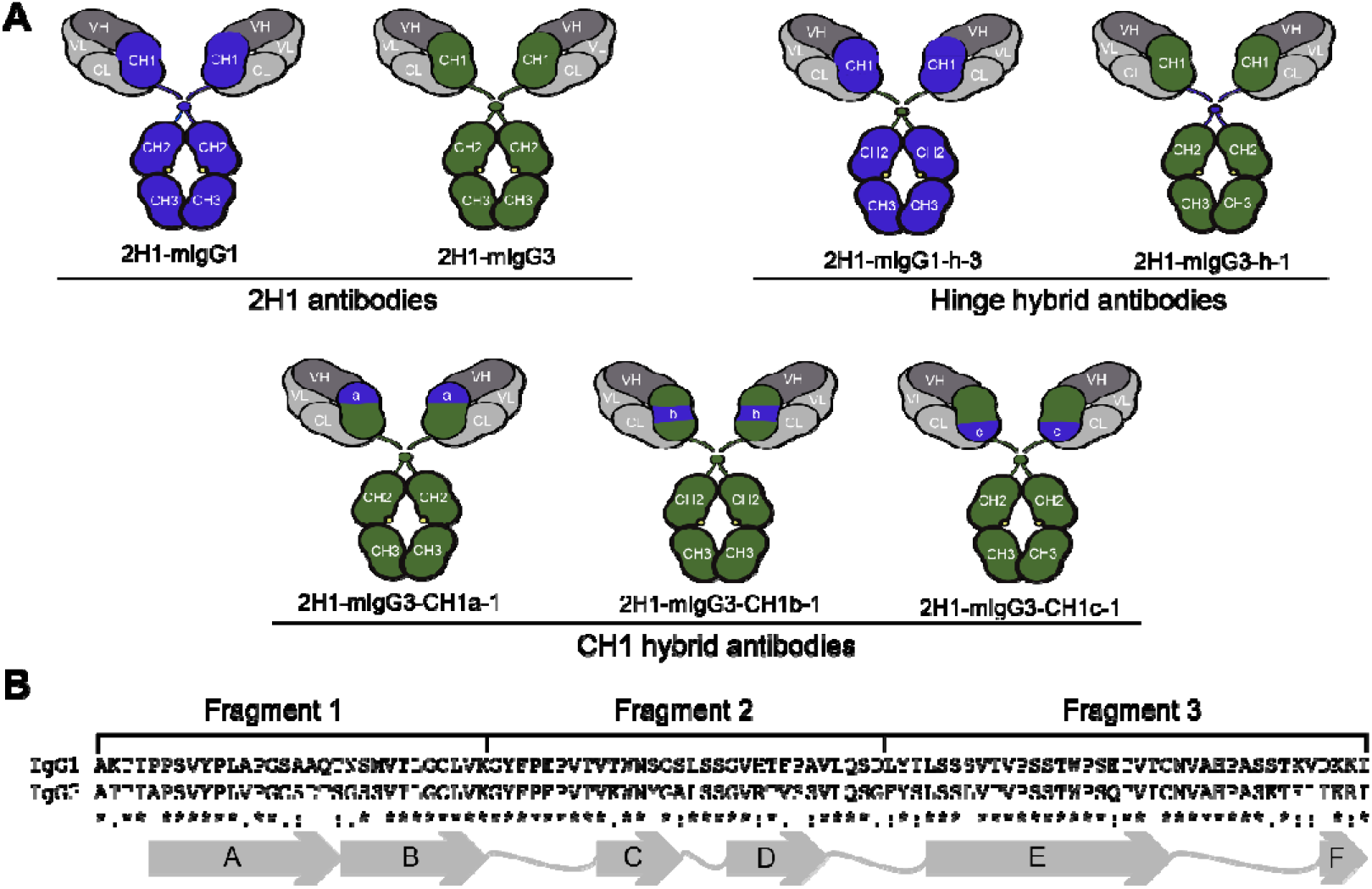
Schematic representation of the recombinant antibodies. (A) 2H1-mIgG1 sequences are represented in blue, whereas 2H1-mIgG3 are represented in green. The three different regions of the CH1 domain that were swapped in the CH1 hybrid antibodies are labeled as “a”, “b” and “c”. (B) Primary and secondary structure of the mIgG1 and mIgG3 CH1 domains, indicating fragments “a”, “b” and “c”.

### Recombinant antibodies - production

CHO dhFr−/−cells were seeded at 5 × 10^4^ cells/mL in 6-well plates 24 h before transfection and then co-transfected with the 2H1-VL containing vector (2H1-VL_pFUSE2-CLIg-mK) plus one of the five hybrid 2H1-VH containing vectors mentioned above, along with the DHFR containing vector (Addgene) to produce 2H1-mIgG3-CH1a-1, 2H1-mIgG3-CH1b-1, 2H1-mIgG3-CH1c-1, 2H1-mIgG1-h-3 and 2H1-mIgG3-h-1 antibodies. Transfections were made using Lipofectamine 2000 (Invitrogen), following manufacturer instructions. Approximately 72 hours post-transfection, we started selection by addition of Zeocin (Invitrogen) at 2 mg/mL, Blasticidin (Gibco) at 10 μg/ml and the cells were kept in negative selection of HT. After two weeks of selection the gene amplification was done using MTX at 10 μM/well, and then in the third week we obtained stable antibody producing cells. The culture volume was then increased to 1 L to produce the first lot of recombinant antibodies. Subsequent lots of the recombinant antibodies were produced by thawing and expanding frozen antibody producing cells.

Additionally, we also produced some of the same recombinant antibodies on a different cell line. NS0 cells were seeded at 8 × 10^5^ cells/mL in 24-well plates 24 h before transfection and then co-transfected with the 2H1-VL_pFUSE2-CLIg-mK vector plus 2H1-VH_pFUSE-CHIg-mG1 or 2H1-VH_pFUSE-CHIg-mG3 vector to produce 2H1-mIgG1 or 2H1-mIgG3 respectively. Seventy-two hours post-transfection, selection of positive transfectants started by addition of Zeocin (Thermo Fisher Scientific) at 1 mg/ml and blasticidin (Thermo Fisher Scientific) at 5 mg/ml. After 3 weeks of selection, stable Ab producing cells were obtained. These cells were then adapted to serum free medium CD Hybridoma AGT (Thermo Fisher Scientific), supplemented with 1x cholesterol (Gibco) and 8 mM L-Glutamine (Sigma Aldrich) and suspension culture at 180 rpm. The recombinant antibodies were produced by thawing and expanding frozen vials of these NSO antibody producing cells.

### Recombinant antibodies - purification and concentration

The purification of 2H1 from NS0 cells consisted of affinity chromatography using rProtein A/Protein G GraviTrap (GE Healthcare). The antibodies were eluted with a 0.1 M glycine-HCL pH 2.7 solution and immediately neutralized with a 1 M Tris-HCl 1 M NaCl pH 9.0 buffer to maintain their stability in solution, then concentrated by ultrafiltration. The 2H1 antibodies from CHO cells supernatant were concentrated in Amicon (Millipore, Danvers, MA) ultrafiltration cells (cutoff 30 kDa).

### Enzyme-linked immunosorbent assay (ELISA)

In solutions with low immunoglobulin amounts the concentration of recombinant antibodies was measured by direct ELISA with antigen. For the antigen-specific ELISA, serial dilutions of mIgG1 (18B7) or mIgG3 (3E5) purified antibodies that had been previously quantified by direct ELISA were used as standards. Plates were coated with 10 μg/mL GXM purified from *C. neoformans* H99 cultures dissolved in PBS. Dilutions of the standards and recombinant antibodies were added and detected with isotype-specific goat anti-mouse polyclonal antibody. For the direct ELISA, plates were coated with serial dilutions of purified myeloma mIgG1/k (MOPC 21) and mIgG3/k (FLOPC 21) standards (Cappel) and dilutions of the recombinant antibodies. After blocking with bovine serum albumin (BSA), the bound antibodies were detected with alkaline phosphatase-conjugated isotype-specific goat anti-mouse polyclonal serum (Southern Biotech).

Competition assays between the 2H1 and 12A1/13F1 were performed by adding a constant amount of one mAb (0,2 to 5 μg/ml) and varying the concentration of another antibody of a different isotype (0 to 100 μg/ml) in an antigen-specific ELISA assay. Ab binding to de-O-acetylated GXM was measured by antigen-specific ELISA, as described above.

### C. neoformans phagocytosis assay

Phagocytosis assays were performed in 96 well tissue-culture treated plates (BD Falcon) containing J774.16 cells plated at 5×10^5^ cell/ml at least 2 h before the experiment. Then, the *C. neoformans* suspension with opsonizing Ab was added at 10 μg/mL, with a macrophage to *C. neoformans* ratio ranging from 1:1 to 1:2. Phagocytosis was allowed to proceed for 2 h at 37°C in 5% CO_2_. Cells were then washed, fixed with methanol at −20°C for 30 minutes and finally stained with Giemsa. Cells were then analyzed under an inverted microscope, counting three fields/well, with at least 100 cells/field. Percent phagocytosis was calculated as the number of macrophages containing one or more internalized *C. neoformans* divided by the total number of macrophages visible in three field. Each experimental condition was done in triplicate.

### Indirect immunofluorescence

The pattern of antibody binding to the *C. neoformans* capsule was evaluated by epifluorescence microscopy. H99 cells were cultivated overnight, washed and diluted to 10^6^ - 10^7^/mL cell density. The cell suspensions were then incubated with antibodies at 10 μg/mL for 1 h at 37°C. After washing with PBS, bound antibodies were detected with an Alexa Fluor^®^ 488-conjugated isotype-specific goat anti-mouse polyclonal serum (*Thermo Fisher Scientific*) for 1 h at 37°C. After washing, stained fungi were mounted on slides with ProLong^®^ Gold Antifade Mounting medium (*ThermoFisher*) and viewed with Zeiss Axio Observer Z1 microscope. For some cells, Z-stacks were collected and subjected to a constrained iterative deconvolution algorithm on Zeiss ZEN software, followed by processing and 3D reconstruction on ImageJ and VOXX2 softwares.

### Statistical analysis

Phagocytosis assays were analyzed with Fisher’s exact test on GraphPad Prism 6,0 software (CA, USA).

## Results

### Production and validation of recombinant antibodies

To understand the structural basis of the differences in mIgG1 and mIgG3 immunofluorescence pattern we had previously observed, we produced more of these two recombinant antibodies (2H1-mIgG1 and 2H1-mIgG3) and five new hybrid antibodies (Figures 1A-B). We hypothesized that the CH1 domain and the hinge are the C region domains most likely to affect the paratope structure and antigen binding, due to the mIgG protein structure (25). Two hybrid antibodies were made to study the role of the hinge domain: 2H1-mIgG1-h-3 is an mIgG1 whose hinge has been substituted by the mIgG3 hinge and 2H1-mIgG3-h-1 is an mIgG3 whose hinge has been swapped with the corresponding mIgG1 sequence. The other three antibodies were used to study the role of the CH1 domain. 2H1-mIgG3-CH1a-1, 2H1-mIgG3-CH1b-1, 2H1-mIgG3-CH1c-1 are all mIgG3 antibodies in which three different portions of the CH1 sequence were swapped with their mIgG1 counterparts. These portions a, b and c correspond to three different regions of the CH1 domain that differ between the isotypes (Figure 1B). Following the IMGT nomenclature, fragment a corresponds to beta-sheets A and B; fragment b includes beta-sheets C and D, loops BC and CD and part of loop DE; fragment c is composed of beta-sheets E and F and the remaining CH1 domain loops.

Following production and concentration, we confirmed that the recombinant antibodies bound to GXM by ELISA (Figure 2A). To determine if the antibodies could bind to the capsule and mediate phagocytosis, we coated *C. neoformans* with 2H1 (mIgG1 and mIgG3) and hybrid 2H1 antibodies (2H1-mIgG3-CH1a-1, 2H1-mIgG3-CH1b-1, 2H1-mIgG1-h-3 and 2H1-mIgG3-h-1). The coated *C. neoformans* cells were then exposed to J774.16 cells, which phagocytosed fungi coated with 2H1-mIgG1 (74%), 2H1-mIgG3 (42%), 2H1-mIgG1-h-3 (40%) and 2H1-mIgG3-h-1 (39%) at a similar rate as the 3E5 hybridoma mAb positive control (Figure 2B-C). The CH1 hybrid recombinant antibodies were also functional, but promoted phagocytosis at a significantly (p<0.0001) lower rate: 2H1-mIgG3-CH1a-1 (10%), 2H1-mIgG3-CH1b-1 (17%). We did not obtain 2H1-mIgG3-CH1c-1 in sufficient concentrations for phagocytosis analysis.

**Figure 2.**
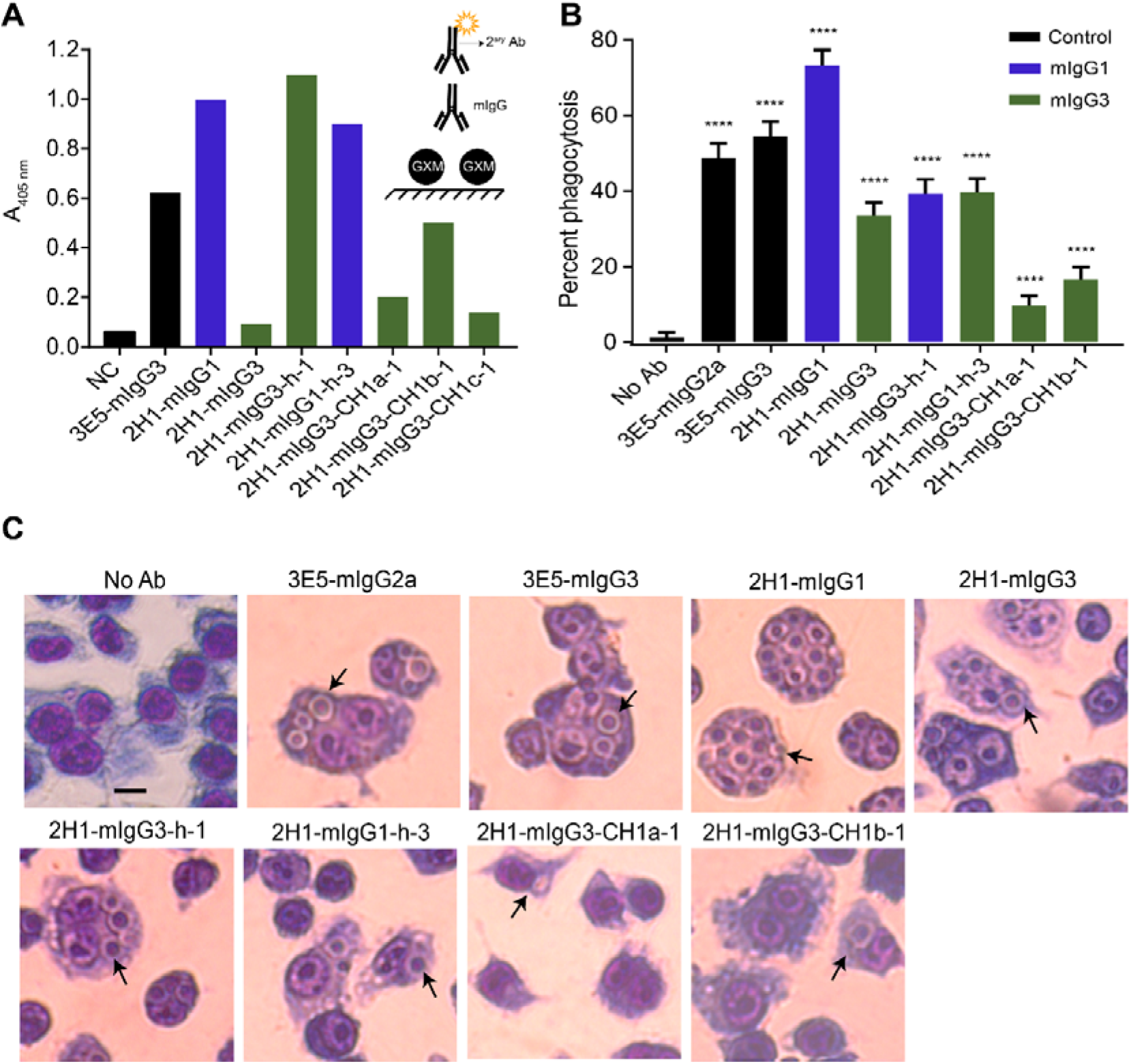
Production and validation of the recombinant antibodies. (A) ELISA experiment with *C. neoformans* capsular polysaccharide and the recombinant antibodies. Bars represent the mean absorbance from a duplicate experiment. (B) Phagocytosis assay with J774.16 cells and *C. neoformans* opsonized with 10 μg/mL of each recombinant antibody. Cells were co-incubated at a 1:2 (macrophage: yeast) ratio, stained and imaged. Bars represent the percentage of macrophages with at least one internalized fungal cell and the 95% confidence interval (Wilson/Brown). **** p<0.0001. (C) Representative images from the phagocytosis assay. Arrows point to macrophages with internalized yeast.

In addition to the recombinant antibodies produced in CHO cells, we produced recombinant 2H1-mIgG1 and 2H1-mIgG3 in the murine myeloma NS0 cells. Both antibodies bound GXM by ELISA (Figure S1A) and successfully mediated phagocytosis (Figure S1B).

### Hinge is necessary but not sufficient to change V binding to antigen

Having validated the serological characteristics of the recombinant antibodies, we next evaluated their binding pattern on the *C. neoformans* capsule. Immunofluorescence staining patterns have been considered good indicators of antibody-mediated protection, such that annular staining correlates with protection in animal models and punctate staining correlates with lack of protection or even disease enhancement (16, 37). As positive controls, we used two hybridoma-derived antibodies, 3E5-mIgG2a and 3E5-mIgG3, known to bind with annular and punctate patterns respectively (17). Our results matched these expected patterns (Figure 3A). The immunofluorescence patterns for 2H1-mIgG1 (annular) and 2H1-mIgG3 (punctate) also matched those previously described for hybridoma-derived 2H1 (37) as well as for the recombinant antibodies and were reproducible with antibodies produced in a different cell line (Figure S1C-D). All three hybrid antibodies in which CH1 fragments were swapped by their mIgG1 counterparts (2H1-mIgG3-CH1a-1, 2H1-mIgG3-CH1b-1 and 2H1-mIgG3-CH1c-1) bound with the punctate pattern expected for mIgG3 antibodies. In contrast, the mIgG3 antibody with an mIgG1 hinge (2H1-mIgG3-h-1) bound with an annular pattern, demonstrating that the hinge is necessary for the punctate binding. The converse antibody, 2H1-mIgG1-h-3, presented the same annular pattern observed for 2H1-mIgG1, which suggests that the mIgG3 hinge is not sufficient for the punctate pattern. To confirm our observations, we collected Z-stacks that were deconvolved and 3D-reconstructed (Figure 3B).

**Figure 3.**
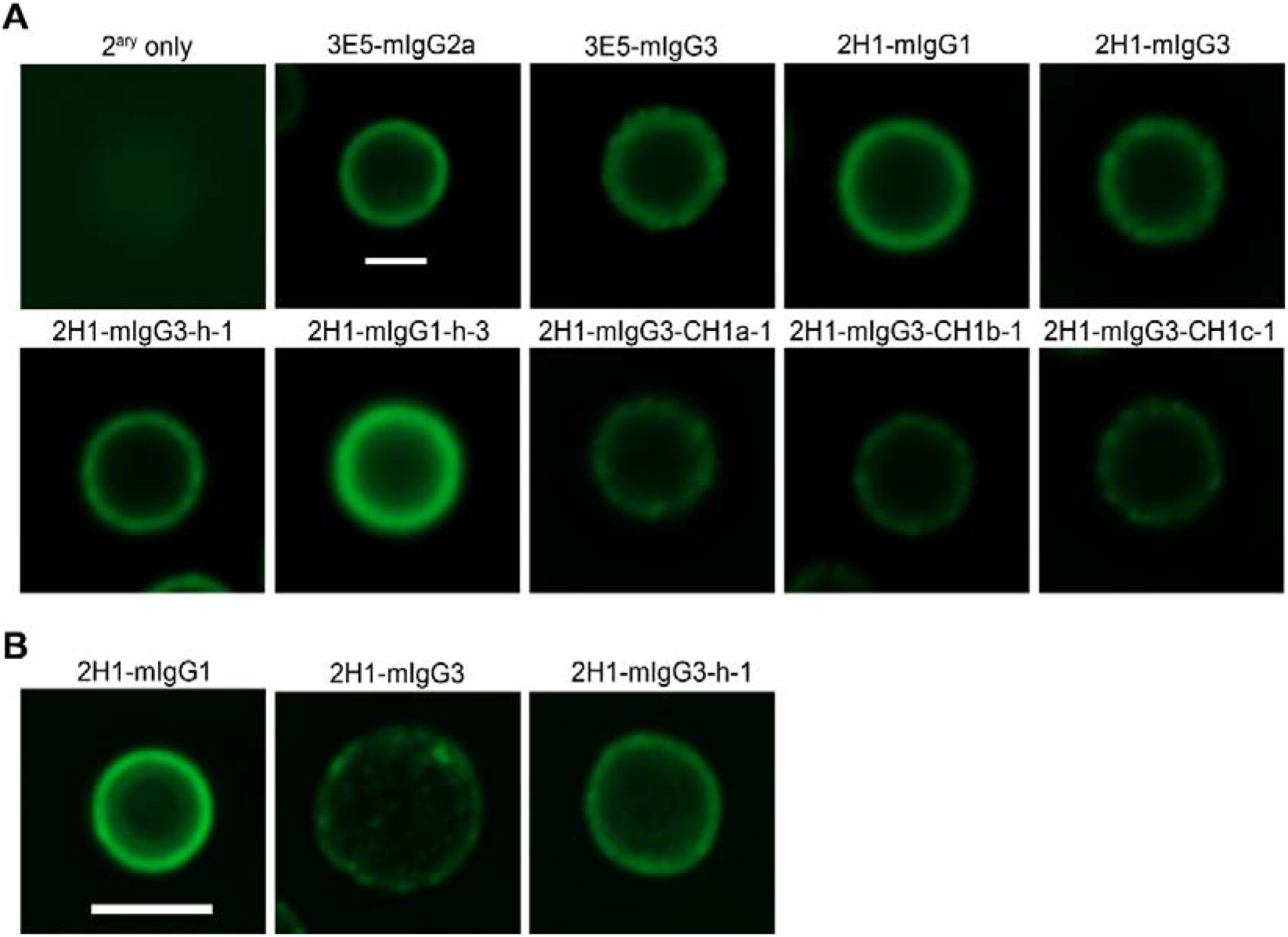
Immunofluorescence pattern of 2H1 antibodies. A) Indirect immunofluorescence of 2H1 antibodies. 2H1-mIgG1 showed annular fluorescence pattern while 2H1-mIgG3 showed punctate pattern, similar to 3E5-mIgG2a and 3e5-mIgG3 patterns respectively. Antibody 2H1-mIgG1-CH1-h-3 showed annular pattern, while antibodies 2H1-mIgG3-CH1a-1, 2H1-mIgG3-CH1b-1 and 2H1-mIgG3-CH1c-1 were punctate. Only 2H1-mIgG3-h-1 changed its punctate pattern to annular. The control, which the fungus was incubated with secondary antibody, did not reveal significant fluorescent signal. The bar scale used in the image was 10 micrometers. (B) 3D representation of 2H1 immunofluorescence pattern. It can be clearly observed that immunofluorescence pattern of 2H1-mIgG3-h-1 was annular, similarly to that of 2H1-mIgG1 and differently to 2H1-mIgG3 punctate pattern.

### 2H1 antibodies bind to a different epitope than 13F1

A competition ELISA assay using GXM with mAbs 2H1, 12A1 and 13F1 showed that 2H1-mIgG1 and mIgG3 antibodies were able to recognize a similar epitope as 18B7 and 12A1 (Figure 4A); but they were not able to recognize the same epitope as 13F1 (Figure 4B).

**Figure 4.**
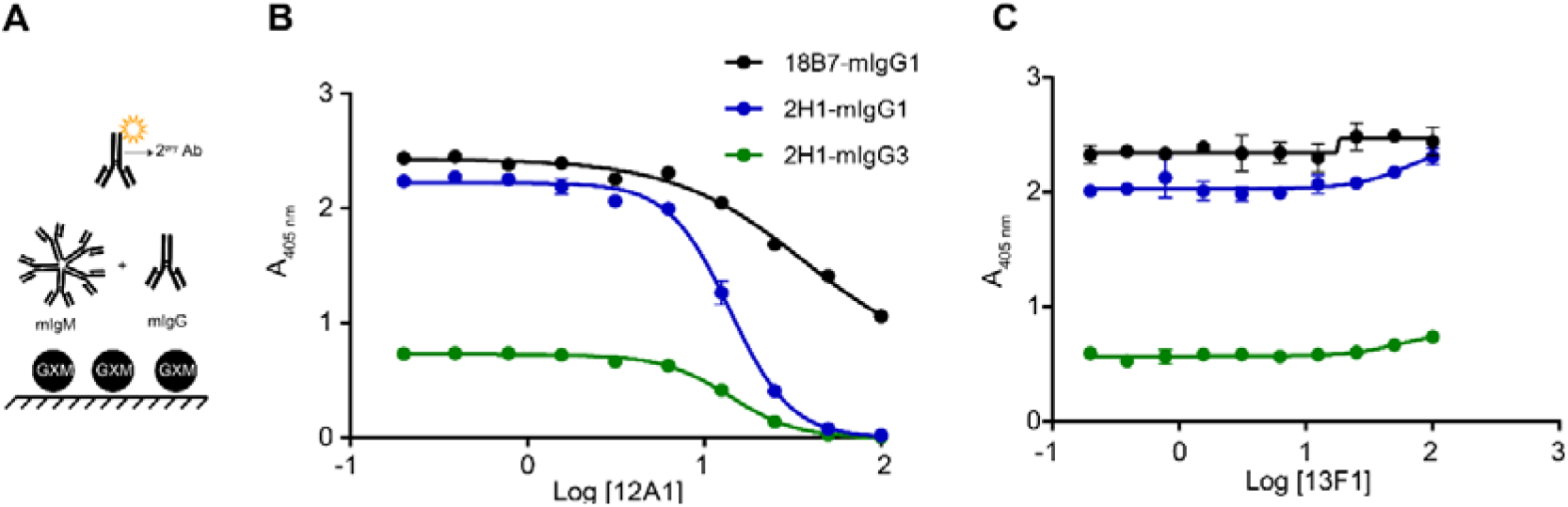
2H1 antibodies binding to GXM epitope. (A) Competitive GXM-ELISA with 2H1 and 12A1 antibodies. 2H1-mIgG1 (blue line), 2H1-mIgG3 (green line) and control antibody (black line) competed for same epitope of 12A1 antibody, an IgM that presents an annular fluorescence pattern. (B) Competitive GXM-ELISA with 2H1 and 13F1 antibodies. 2H1-mIgG1, 2H1-mIgG3 and control antibody did not compete for the same epitope of 13F1, an IgM that presents a punctate fluorescence pattern. The scheme in the figure represents the ELISA assay conditions. The graphs represent mean absorbance of duplicate experiment.

### 2H1-mIgG3 binding to GXM is different than 2H1-mIgG1 binding

To better understand the differences between 2H1-mIgG1 and 2H1-mIgG3 antigen binding, we tested their binding to native and de-O-acetylated EPS obtained from *cas1Δ* or chemical reaction (Figure 5A). We found that the relative strength of 2H1 binding to de-O-acetylated GXM was mIgG3 > mIgG1 (Figure 5B). Consistent with this observation the 3E5 reactivity to de-O-acetylated GXM was similar (17). The antibody reactivity results reinforce immunofluorescence patterns found to 2H1, 3E5 and 18B7. Because the switch variants have identical variable region the stronger binding by mIgG3 isotypes suggests that C region may affect the V region specificity.

**Figure 5.**
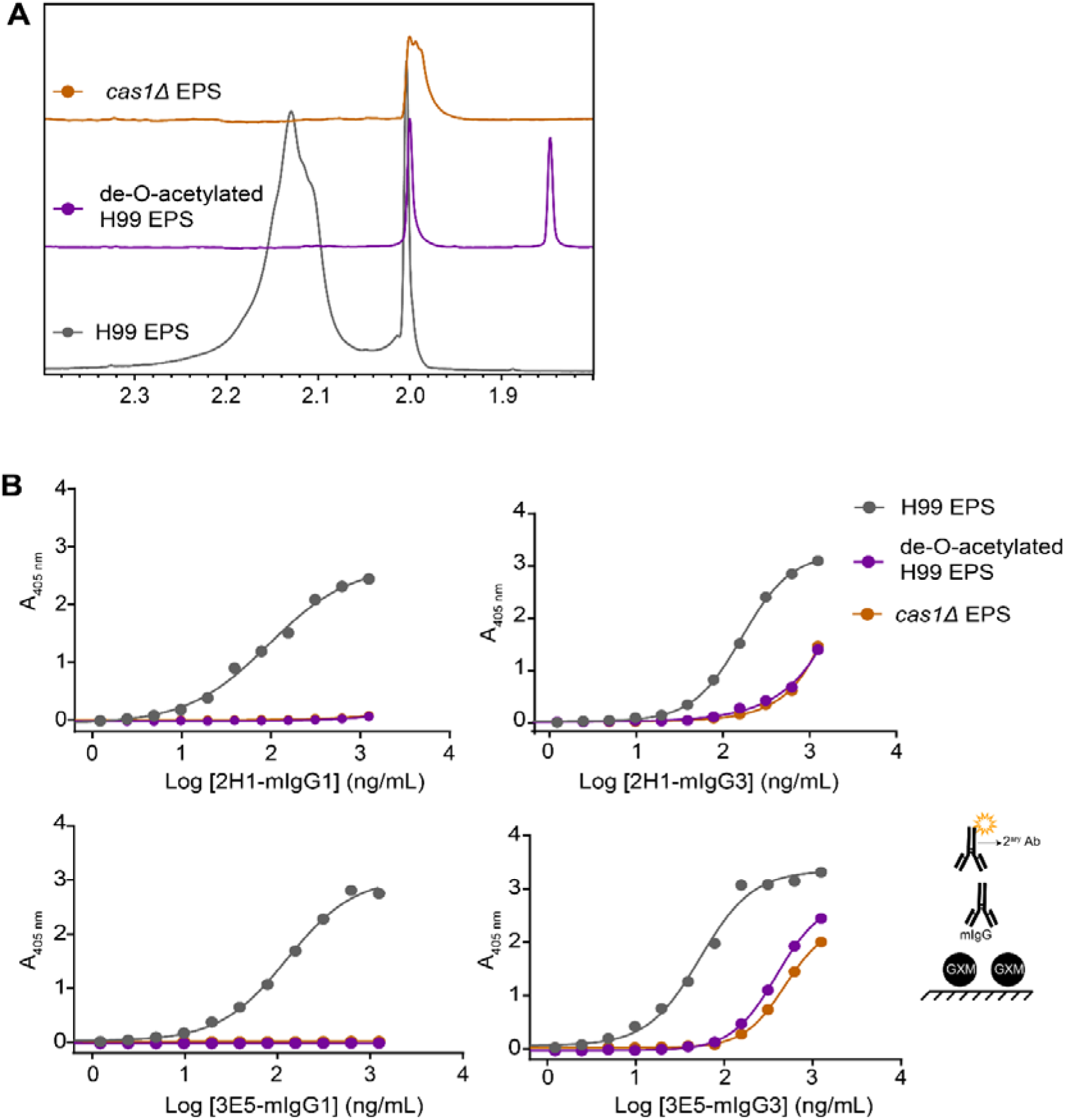
2H1 antibodies binding to de-O-acetylated GXM. (A) Analysis of EPS de-O-acetylation by NMR. The peak in range 2.0 represents de-O-acetylation fraction of *cas1*Δ (orange line), chemically modified EPS (purple line) and non-de-O-acetylated EPS (gray line). (B) Interaction ELISA of 2H1 antibodies to de-O-acetylated GXM. 2H1-mIgG3 antibody binds to de-O-acetylated GXM, while 2H1-mIgG1 antibody does not bind. The graphs represent the mean absorbance of three technical replicate.

## Discussion

Among the immunoglobulin regions, it was previously believed that the variable was the only one that influenced binding of antibody to antigen (12, 13). After three decades of study, this definition has been updated to include the constant region which was shown to influence the functionality of the paratope by conformational change on V region (26), influence the chemical and electronic environment (27) and alter of the multivalent interactions (38). Immunofluorescence studies carried out with mIgG isotypes of identical variable region against the *C. neoformans* capsule were important to reveal differences in antibody binding under the influence of the constant region. However, only three different sets of IgG antibodies (3E5, 4H3 and chimeric 18B7) (16, 17) were used to characterize the binding pattern to the fungus, being punctate when the opsonizing antibody is a mIgG3 isotype and annular when it is of another isotype. In this way, we created another set of recombinant mIgG, the 2H1-mIgG1 and mIgG3, to reinforce that the binding patterns do not occur by chance. Interestingly, the 2H1 antibodies reproduced the immunofluorescence results, where mIgG3 showed a punctate binding pattern, whereas mIgG1 showed an annular pattern when bound to GXM (39).

Differences in the way IgG isotypes with identical V region sequences bind to antigen are related to the recognition of different epitopes (36). Therefore, we performed ELISA with different forms of GXM (O-acetylated and de-O-acetylated) to evaluate the binding of the different 2H1 isotypes to these antigens. What is observed in the literature is that the mIgG1 isotype binds to O-acetylated GXM, but does not bind to de-O-acetylated GXM; and that mIgG3 binds to both acetylated and de-o-acetylated GXM (36). We obtained the same results when we evaluated the 2H1-mIgG1 and mIgG3 binding to antigen and this led us to the hypothesis that the annular and punctate binding observed in immunofluorescence occurs due to isotypes recognizing different epitopes. Competition ELISA performed between 2H1-mIgG1/mIgG3 antibodies and 12A1/13F1 antibodies also revealed that the mIgG1 and mIgG3 antibodies compete for the 12A1 epitope in greater and lesser affinity, respectively, even though they have the same variable region. Because 12A1 and 2H1 use the same VH (IGHV5-6-2) and JH (IGHJ2) genes and they are classified as members of class II mAbs (21), the competition for the same epitope was expected (17). Although 13F1 also belongs to the same family, its VH mutations (28) can reveal residues important in binding to GXM that probably affected the competition with the other antibodies. As for the 2H1-mIgG3 competition result, it was found that 3E5-mIgG3 - the most 2H1 identical antibody - competed with 12A1 but not with 13F1 (28).

Although several studies have been carried out, such as those mentioned above, the mechanism by which the constant chain influences the binding of antibodies has not yet been identified. Molecular modeling based on crystallographic data of the VH and CH1 of antibodies 3E5 mIgG3 and 2H1 mIgG1, suggested that the loop between VH and CH1 may exert structural or kinetic influence on the paratope (40, 41). In addition, homology modeling analysis of the IgG subclasses identified three regions with structural differences in the CH1 domain that can form structural isomers and can change the kinetics of the antigen-antibody complex (14). In order to investigate whether CH1 influences VH we created three antibodies with changes in the CH1 region, but we were unable to observe direct influence of the constant region on binding to *C. neoformans*. Biophysical studies suggest that functional change of the paratope may not occur due to a single domain, but rather due to the influence of a set of regions (42). This hypothesis may also explain why we observed the influence of the hinge region on antigen binding with the 2H1-mIgG3-h1 hybrid, but not with the 2H1-mIgG1-h3. Thus, we propose that the hinge region is necessary to influence the antigen binding, but it is insufficient to do this alone. When the hinge of 2H1-mIgG3 was replaced by that of mIgG1, there was a change in the pattern of binding to GXM, where it ceased to be punctiform and became annular. Since the primary, secondary structure (26) and the degree of Fab-Fc flexibility of the hinge region differs between the IgG isotypes, this suggests that this region is important for spatial reach and antigen binding, as demonstrated by the small-angle X-ray scattering experiment and modeling performed with 3E5 antibodies (25), the ELISA experiments with chimeric antibodies (38) and also the alignment of amino acids performed in this study (Fig. 1).

The experimental evidence presented herein lead us to two conclusions: 1) the hinge region is important, but not the only factor influencing antibody binding to antigen; 2) The constant region affects the variable region, impacting the way antibody binds to the antigen, changing its specificity and affinity to the epitope and this event does not happen by chance. Consequently, antibody molecules produced against the same antigen may or may not allow V-C allosteric effects, thus making the immunoglobulin function unpredictable. This unpredictability can have a negative impact on antibody-based therapies, since it is not possible to predict if the result of the antigen-antibody interaction will be effective, especially in the case of active therapies where antibodies are produced *in vivo* in a complex microenvironment. We note that different V-C combinations differ on the ability of the C region to affect specificity and that somatic mutations could affect the permissiveness of this phenomenon, thus adding more unpredictability to properties of assembled immunoglobulins (12). For example, monoclonal mIgG antibodies against the same antigenic determinant, but of different isotypes, show different effectiveness in treating disease – some isotypes being protective whereas others are non-protective or even have the potential to aggravate the disease (22, 43–45). This problem is also noticed even in passive therapies with commercially available antibodies (46).

Our results underscore the need to consider the immunoglobulin molecule holistically with regards to its interactions with antigen rather than focusing only on V region domains. Therefore, the results of this work reinforce the importance of understanding the intra and intermolecular cooperativity mechanism of IgG domains, in order to contribute to the safety and efficacy of antibodies during the engineering and therapeutic processes.

## Acknowledgements

We would like to acknowledge Dr. Ildinete Silva Pereira from the University of Brasilia, the Genomic Sciences and Biotechnology facility at Catholic University of Brasilia and the Immunology Core at Johns Hopkins University for the technical support.

## Abbreviations

Ab: Antibody
C: constant
CH: constant heavy chain
dhFr: dihydrofolate reductase
CL: constant light chain
DME: Dulbecco’s Modified Eagle’s
ELISA: Enzyme-Linked Immunosorbent Assay
Fab: Fragment antigen-binding
FITC: fluorescein isothiocyanate
GXM: Glucuronoxylomannan
IgG: immunoglobulin γ
mIgG: murine immunoglobulin γ
IgM: immunoglobulin μ
mIgM: murine immunoglobulin μ
mAbs: monoclonal antibodies
mIgG: murine immunoglobulin gamma
SAXS: Small-angle X-ray scattering
sRBC: Sheep red blood cells
VH: variable heavy chain
VL: variable light chain

