## Supplemental figure 1 for "Hinge Influences in Murine IgG Binding to *Cryptococcus neoformans* Capsule"

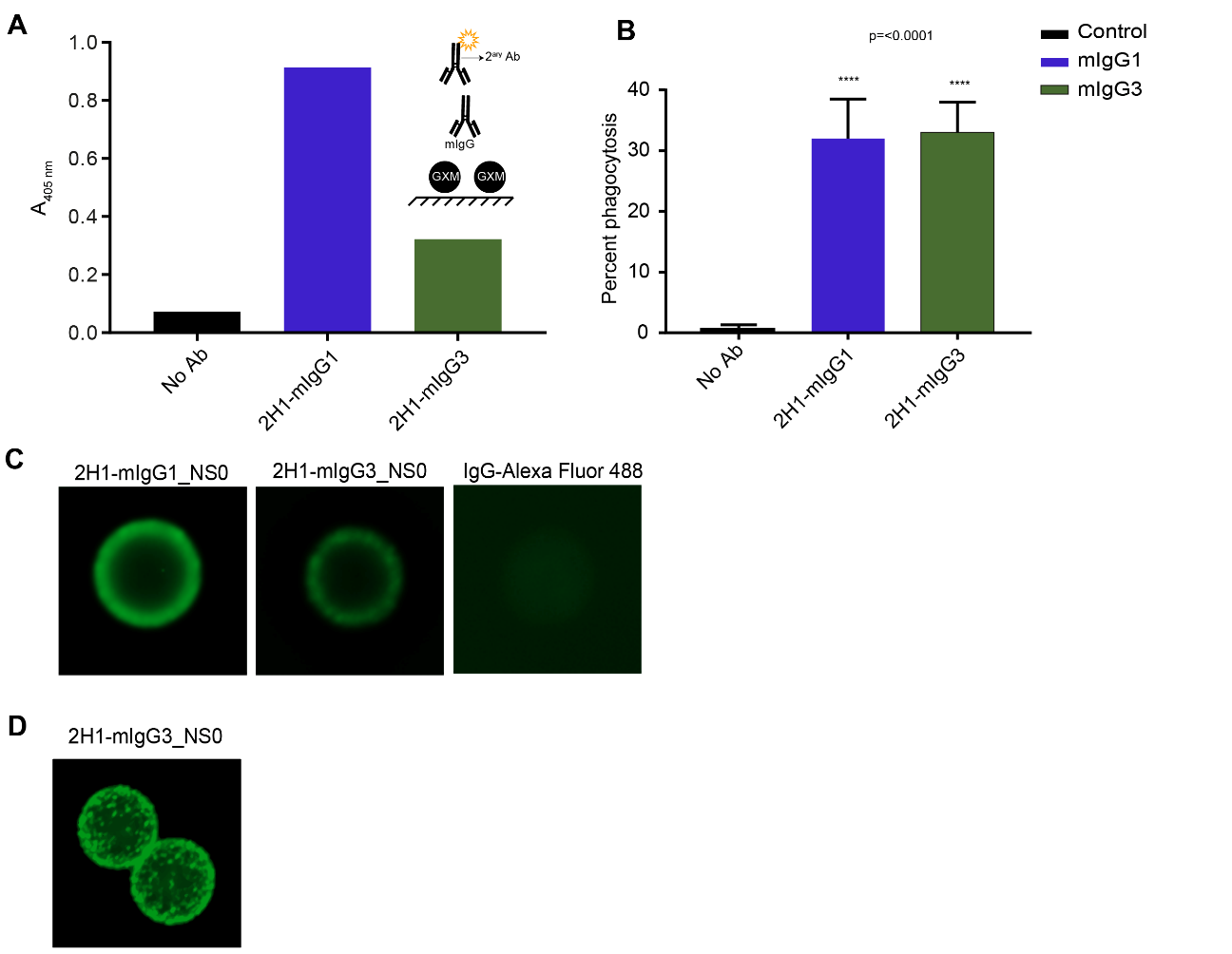

*Supplemental material – 2H1 antibodies from NS0 cell*

(A) ELISA experiment with *C. neoformans* capsular polysaccharide and the recombinant antibodies. Bars represent the mean absorbance from a duplicate experiment. (B) Phagocytosis assay with J774.16 cells and *C. neoformans* opsonized with 10 µg/mL of each recombinant antibody. Cells were co-incubated at a 1:2 (macrophage: yeast) ratio, stained and imaged. Bars represent the percentage of macrophages with at least one internalized fungal cell and the 95% confidence interval (Wilson/Brown). **** p<0.0001. (C) Representative images from indirect immunofluorescence assay. (D) 3D reconstruction of 2H1-mIgG3_NS0 immunofluorescence pattern.
